# Frontotemporal Coordination Predicts Working Memory Performance and its Local Neural Signatures

**DOI:** 10.1101/2020.03.05.976928

**Authors:** Ehsan Rezayat, Mohmmad-Reza A. Dehaqani, Kelsey Clark, Zahra Bahmani, Tirin Moore, Behrad Noudoost

**Affiliations:** School of Cognitive Sciences, Institute for Research in Fundamental Sciences (IPM), PO Box 1954851167, Tehran, Iran; Cognitive Systems Laboratory, Control and Intelligent Processing Center of Excellence (CIPCE), School of Electrical and Computer Engineering, College of Engineering, University of Tehran, Tehran, Iran; Department of Ophthalmology and Visual Sciences; University of Utah; Salt Lake city, UT; Department of Biomedical Engineering, Tarbiat Modares University, Tehran, Iran; Department of Neurobiology Stanford University, Stanford, California 94305, USA; Howard Hughes Medical Institute, Stanford University, Stanford, California 94305, USA

## Abstract

Neurons in some sensory areas reflect the content of working memory (WM) in their spiking activity. However, this spiking activity is seldom related to behavioral performance. We studied the responses of inferotemporal (IT) neurons, which exhibit object-selective activity, along with Frontal Eye Field (FEF) neurons, which exhibit spatially-selective activity, during the delay period of an object WM task. Unlike the spiking activity and local field potentials (LFPs) within these areas, which were poor predictors of behavioral performance, the phase-locking of IT spikes and LFPs with the beta band of FEF LFPs robustly predicted successful WM maintenance. In addition, IT neurons exhibited greater object-selective persistent activity when their spikes were locked to the phase of FEF LFPs. These results demonstrate a key role of coordination between prefrontal and temporal cortex in the successful maintenance of visual information during WM.

## Introduction

High level brain areas such as prefrontal cortex (PFC) and parietal areas are believed to be central in the control and execution of behavioral plans^1^. How these areas interact with sensory areas to maintain task-relevant information is not well understood. Neurons in prefrontal cortex exhibit sustained spiking activity that encodes the content of WM^2^. However, since feature selectivity in some PFC areas seems insufficient to match the resolution of WM, a *sensory recruitment* model of WM has been suggested, in which brain areas involved in sensory processing also contribute to memory maintenance^3,4^. Such models involve interactions between PFC and sensory areas during memory maintenance. Variations of this idea range from a more modular perspective, in which specific portions of PFC exhibiting memory activity send that signal to sensory areas^5,6^, or a split of abstract vs. detailed information^7^ (perhaps varying based on task demands^8^), to more distributed versions emphasizing the content-specific communication between areas rather than spiking activity within either^9^. These conceptual models are not necessarily mutually exclusive, but they all predict that memory-related activity in sensory areas should be related to WM performance, a prediction that is discrepant with much of the existing neural data. Studies have shown that even when the content of WM is present in spiking activity within sensory areas, it is either only weakly correlated^10,11^ or not correlated at all^12,13^ with behavioral performance, casting doubt on its role in memory maintenance.

In order to examine how the PFC interacts with sensory areas in support of WM maintenance, we simultaneously recorded spiking activity and LFPs within the FEF and IT in monkeys performing an object WM task. We sought to determine which aspects of the neural response were most closely linked to successful WM maintenance. We found that neither the spiking activity nor the LFP power spectrum within either of these areas was a strong predictor of the animal’s performance on this task. However, the synchronization between the two areas (phase-phase locking), particularly in the beta band, was a key predictor of successful object memory maintenance. Even though the object selectivity within IT did not differ between correct and wrong trials, the timing of spikes in IT were coordinated with the phase of the FEF LFP, and when the two areas were more strongly locked, IT spiking activity showed greater selectivity for the object held in memory. Thus, beta band synchrony between IT and FEF predicted both behavioral performance and the strength of object-selective persistent activity in IT.

## Results

We recorded spiking activity and LFP signals simultaneously from the FEF and IT cortex of two monkeys (M1, M2) using two single electrodes (228 IT units, 58 IT LFP sites, 161 FEF units, 86 FEF LFP sites). To examine the neural basis of object WM, we trained two monkeys to perform an object delayed-match-to-sample (DMS) task (Fig. 1a) in which they had to remember the identity of a sample stimulus throughout a delay period (1 second). The sample stimulus appeared either within the FEF RF (In condition) or 180 degrees away (Out condition). Following the delay period, two stimuli appeared in locations rotated 90 degrees relative to the sample location, and monkeys were rewarded for making a saccadic eye movement to the one that matched the sample stimulus, regardless of its location. Different sample objects evoked a comparatively stronger or weaker response in the IT unit being recorded, which was used to define the preferred (Pref) and nonpreferred (NPref) stimulus conditions. Performance of the two animals is shown in figure 1b (correct trials = 70 ± 1% M1, 77 ± 1% M2). For analyses involving the FEF alone, we considered the In condition unless otherwise noted; for analyses involving IT alone, we considered the Pref condition unless otherwise noted. For analyses involving both FEF and IT, we considered the Pref, In condition unless otherwise specified.

**Figure 1.**
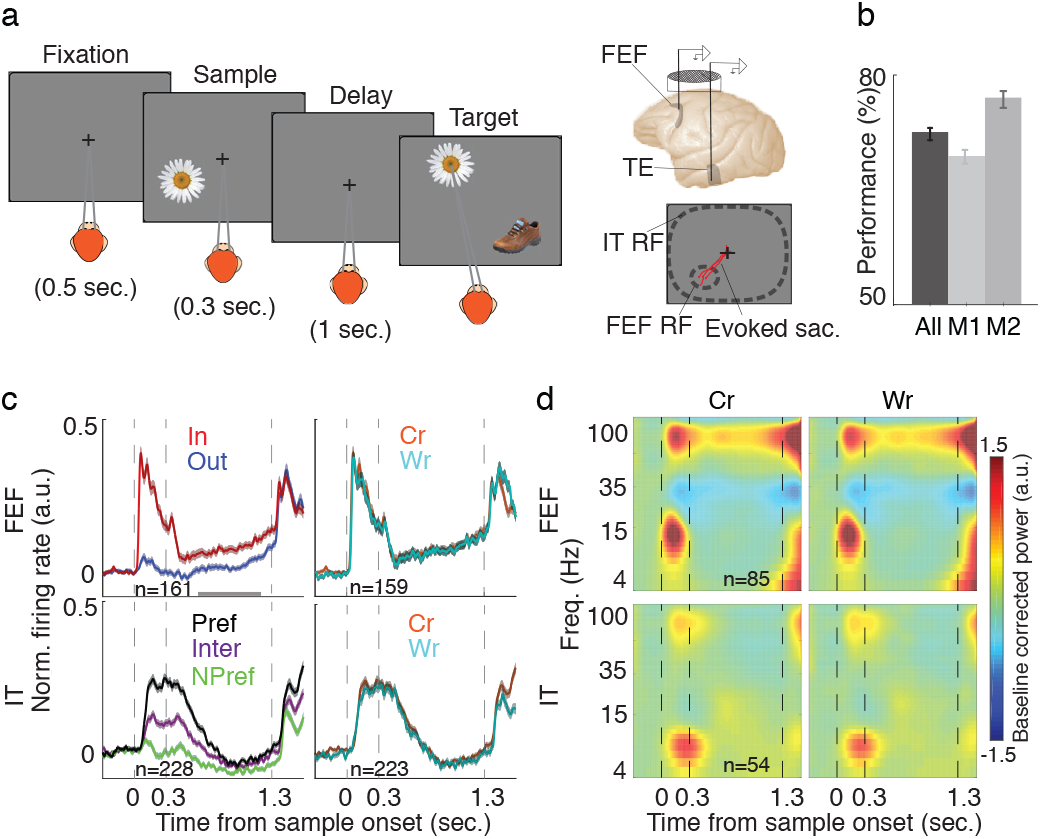
Firing rate and LFP power in IT and FEF were largely unrelated to DMS performance. **a,** Schematic of the object DMS task. Left) After the monkey fixated, and a sample stimulus appeared either in or opposite the FEF neuron’s RF location. The sample stimulus disappeared, and the monkey maintained fixation throughout a blank delay period. The same stimulus appeared again (target) along with a distracter and the monkey needed to saccade to the target stimulus to receive a reward. Right) Recordings were made simultaneously from the FEF and the TE part of IT in single recording chamber. The FEF RF was estimated based on microstimulation-evoked saccades (red lines). The IT RF generally encompassed both sample locations. b, Mean performance of individual monkeys (M1, M2) and their average (All) on the DMS task; error bars show standard error across sessions. **c,** Normalized firing rates of FEF (top) and IT populations (bottom) across the course of the DMS task, when the sample was inside (red) or outside (blue) of the neurons’ RFs, and when the sample object was preferred (black), non-preferred (green), or intermediate (violet) are shown on the left. Right) Normalized firing rate of the FEF (top) and IT populations (bottom) for correct vs. wrong trials. Data (mean ± SE) were smoothed within a window of 10 ms. **d,** Time-frequency maps of normalized LFP spectral power for FEF (top) and IT (bottom) for correct (left) and wrong (right) trials.

### Within-area neural activity does not predict behavioral performance

We first examined whether changes in the FEF or IT firing rate were correlated with behavioral performance on the object WM task. Consistent with an earlier study^14^, the population of FEF units (102 M1, 59 M2) exhibited persistent activity, measured as an increase in normalized firing rate (NFR) during the delay period compared to their baseline (ΔNFR = 0.078 ± 0.011, p < 10^-12^, n = 161; Fig. 1c, upper left). This persistent activity was significant in 68% of FEF units (110/161). This increase in delay-period FEF spiking activity was only slightly greater during correct (Cr) trials than wrong (Wr) trials (ΔNFR = 0.006 ± 0.003, p = 0.041, n = 159; Fig. 1c, upper right). In the population of IT units (170 M1, 58 M2), although 28% of IT units (64/228) exhibited significantly elevated delay period spiking activity vs. baseline, in the whole population the increase in delay period firing rate was not significant (ΔNFR = – 0.004 ± 0.006, p = 0.930, n = 228). Moreover, there was no significant difference in the delay period spiking between the correct and wrong trials (ΔNFR = 0.007 ± 0.005, p = 0.282, n = 223; Fig. 1c, lower right). Delay period activity in FEF and IT was selective for sample location (In-Out) and object identity (Pref-NPref), respectively (FEF spatial selectivity = 0.060 ± 0.012, p < 10^-7^, n = 161; IT object selectivity = 0.049 ± 0.005, p < 10^-15^, n = 228). However, the magnitude of this selectivity did not vary with performance (ΔFEF spatial selectivity = −0.002 ± 0.006, p = 0.415, n = 159; ΔIT object selectivity = – 0.001 ± 0.007, p = 0.459, n = 223). Thus, neither FEF’s nor IT’s average spiking activity was a strong predictor of object WM performance.

Next, we investigated the strength of neural oscillations within each area. We calculated a time-frequency map of the LFP power spectra for each of the FEF sites (51 M1, 35 M2) and IT sites (36 M1, 22 M2)using a wavelet transform (see Methods). During the delay period, FEF power in the beta (β, 20-28 Hz) and gamma (γ, 50-130 Hz) bands reflected the sample location (see Table S1 and Supplementary Information). Despite this spatial selectivity, there was no significant difference in LFP power between correct and wrong trials during the delay period for either the beta or the gamma band (Δpower_β_ = −0.030 ± 0.029, p = 0.078; Δpower_γ_ = −0.011 ± 0.033, p = 0.416, n = 85 sites; Fig. 1d, top). Within IT, there was an increase in alpha band power (α, 8-15 Hz) during the delay period which reflected sample identity (see Table S1 and Supplementary Information). However, as in the FEF, there was no significant difference in alpha band LFP power between correct and wrong trials during the delay (Δpower = 0.009 ± 0.037, p = 0.990, n = 54 sites; Fig. 1d, bottom). Nor did power in any other frequency band reflect performance (Table S1). Overall, although FEF and IT firing rates and LFP power during the delay period were modulated by the sample object and location, none of these measures were strongly predictive of behavioral performance.

### Inter-areal beta band coupling predicts performance and reflects the content of WM

Since neither spiking nor local field activity within FEF or IT was strongly predictive of performance, we next investigated the synchronization between FEF and IT sites in our search for behavioral correlates of WM. As a measure of inter-areal coupling, we calculated phase-phase locking (PPL) between simultaneously recorded LFPs from FEF and IT (36 M1, 22 M2); PPL was calculated across time and frequency, and shuffle corrected (see Methods). Importantly, we observed a significant enhancement in beta band PPL during the delay period compared to baseline (ΔPPL = 0.295 ±0.095, p = 0.005, n = 58), indicating a more consistent phase relationship between beta LFPs in FEF and IT during memory maintenance. A more modest increase in phase locking between areas occurred during the sample period compared to baseline (ΔPPL = 0.209 ±0.098, p = 0.034, n = 58). There was no delay-period increase in PPL in other frequency bands (Table S1). We compared baseline-adjusted delay period PPL for correct vs. wrong trials, and found that in the beta band, PPL was greater for correct trials (ΔPPL = 0.442 ± 0.131, p = 0.002, n = 50; Fig. 2a-c). This enhanced beta band PPL for correct trials was significant after trial matching or without the shuffle correction (Fig. S1). Successful WM performance was thus more strongly associated with the phase locking between the FEF and IT than with the average spiking activity or the power of LFPs within each area.

**Figure 2.**
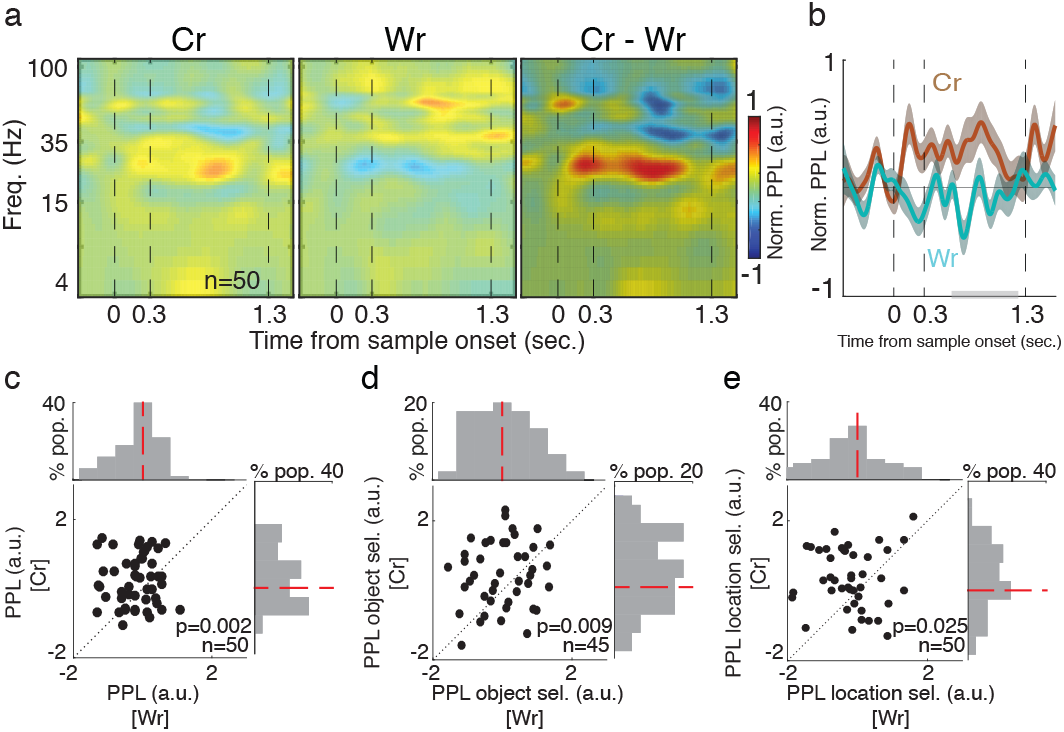
Interareal beta coupling during correct and wrong trials. **a,** PPL between FEF and IT was Ś different for Cr and Wr trials. The timefrequency map of PPL, normalized to baseline across the course of the DMS task for correct (left), wrong (middle), and the difference between Cr and Wr trials (right). **b,** Beta band PPL during β sample and delay differed between Cr and Wr trials. Plot shows the average of beta band PPL across the course of the DMS task for the correct (red) and wrong (blue) trials. Shading shows the SE across recorded pairs. Gray bar indicates the analysis window for (c-e). **c,** Comparison of beta band PPL during the delay period for correct versus wrong trials. Beta band PPL was significantly higher for correct trials. **d,** Comparison of object selectivity of beta band PPL during the delay period for correct versus wrong trials. Object selectivity of beta band PPL was significantly higher for correct trials. **e,** Comparison of spatial selectivity of beta band PPL during the delay period of correct versus wrong trials. Spatial selectivity of beta band PPL was significantly higher for correct trials.

Having identified the beta coupling between areas as a predictor of successful WM maintenance, we next examined whether this measure also related to the content of WM. To do this, we compared the beta band PPL between areas for different sample objects and locations. Measuring the object selectivity (Pref, In – NPref, In), we found that the beta band PPL between the FEF and IT carried information about the identity of remembered objects (object selectivity = 0.520 ± 0.142, p = 0.001, n = 51). No other frequency band showed similar significant object selectivity of the PPL (Table S1, Fig. S1). Moreover, the object selectivity of the beta band PPL was greater during correct compared to wrong trials (Δobject selectivity = 0.513 ± 0.170, p = 0.009, n = 45; Fig. 2d). Thus, the content of object WM was reflected in the beta band PPL between the FEF and IT, and the strength of this encoding was correlated with behavioral performance. We also investigated whether beta coupling encodes the sample location during the delay period by measuring its spatial selectivity (Pref, In – Pref, Out). We found that the beta band PPL during the delay period also carried spatial information (spatial selectivity = 0.364 ± 0.150, p = 0.019, n = 58). In addition, the spatial selectivity of the beta band PPL was greater for correct compared to wrong trials (Δspatial selectivity = 0.474 ± 0.174, p = 0.025, n = 50; Fig. 2e). Thus, beta coupling between the FEF and IT reflected both the sample identity and location of the remembered stimulus, and both the spatial and object selectivity were stronger on correct trials.

In order to investigate the directionality of fronto-temporal coordination, we next examined the phase difference between FEF and IT LFPs. On correct trials, we observed a positive phase difference between FEF and IT, indicating that FEF led IT. Figure 3a shows a distribution of phase differences between FEF and IT for correct (top) and wrong (bottom) trials. On correct trials, the FEF beta phase led IT (μ = 19.309°, κ = 1.400, p < 10^-7^, n = 50, Von Mises test) and for wrong trials IT led FEF (μ = −14.500°, κ = 0.857, p < 10^-4^, n = 50, Von Mises test). The variability in phase difference values was also reduced on correct trials (k_Cr_ = 1.463 ± 0.354, κ_Wr_ = 0.904 ± 0.198, p = 0.007, n = 1000x bootstrap). These results suggest that the FEF plays a leading role in the beta band coupling during correct trials.

**Figure 3.**
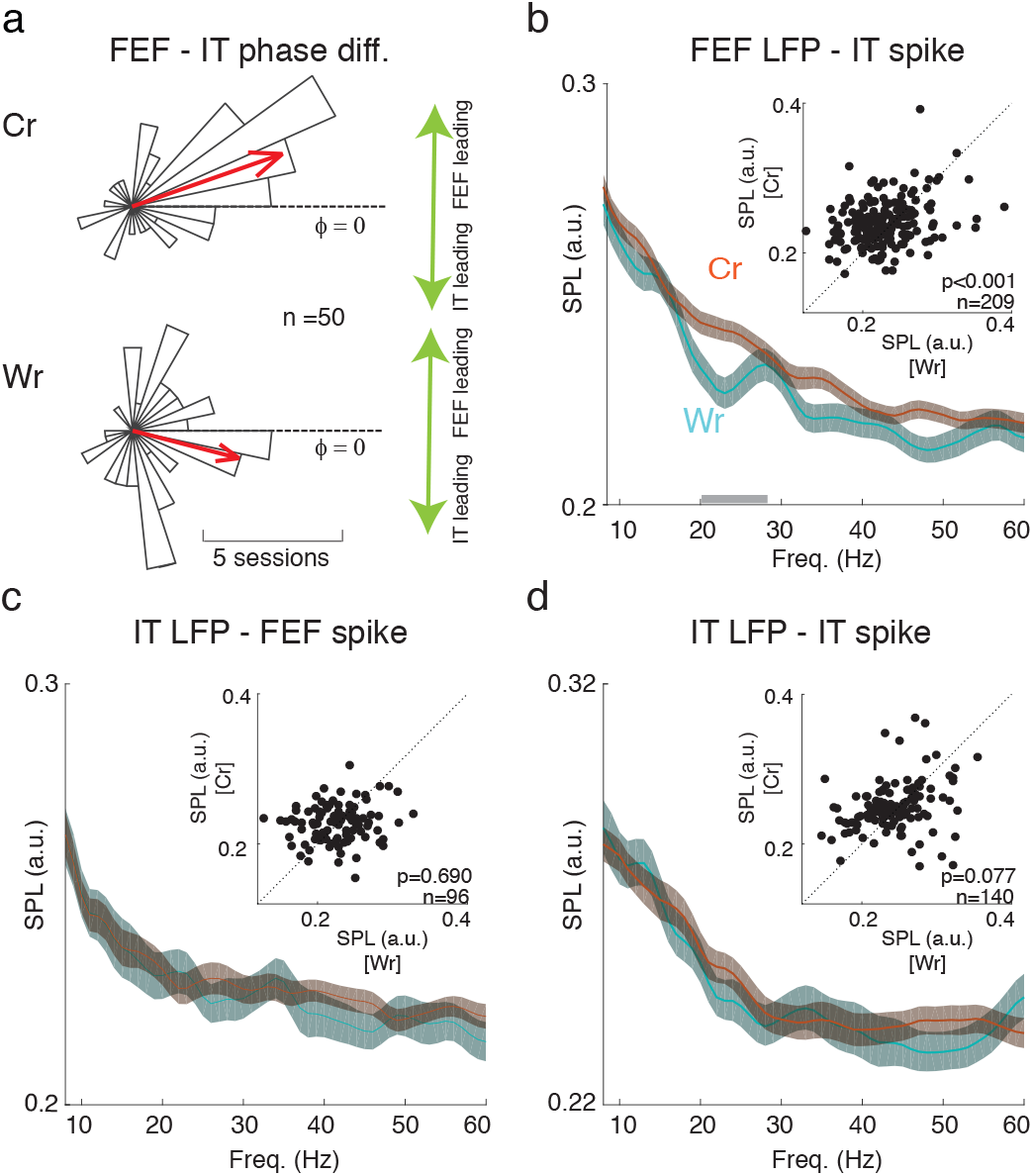
Inter-areal beta coupling, led by FEF, reflected performance. **a,** Beta band phase differences between areas reflected performance. The polar plot shows a histogram of phase differences between the FEF and IT LFPs in the beta band for correct trials (top) and wrong trials (bottom). Arrows show the circular means across all pairs. **b,** Beta band SPL between IT spikes and FEF LFPs reflected performance. Plots show the SPL for correct (red) and wrong (blue) trials across frequencies for all pairs of IT units and simultaneously recorded FEF LFPs. Gray bar on x-axis indicates the beta frequency range. Data were smoothed within a window of 1 Hz and represented as mean ± SE. Inset scatter plots compare beta range SPL for correct vs. wrong trials. **c-d**, Plots show the SPL for correct and wrong trials across frequencies for all pairs of FEF units and simultaneously recorded IT LFPs (c), and for IT units and simultaneously recorded IT LFPs (d). Smoothing and scatter plot as **in** (b).

Next, we analyzed whether the increase in beta band PPL was accompanied by an increase in inter-areal spike-phase locking (SPL). To calculate SPL, each spike was assigned a phase value based on the simultaneously recorded LFP; SPL was the circular average of all spike phases at a particular frequency (see Methods). We calculated SPL for both FEF spikes and IT LFPs, and IT spikes with FEF LFPs, but only the latter showed significant correlations with behavior (Fig. 3b-d). Figure 3b illustrates SPL between FEF LFPs and IT spikes for correct vs. wrong trials across various frequencies; there was an increase in the theta (θ; 4-8 Hz), beta, and gamma band SPL on correct compared to wrong trials (ΔSPL_θ_ = 0.007 ± 0.004, p = 0.013; ΔSPU_β_ = 0.010 ± 0.003, p < 10^-3^; ΔSPU_γ_ = 0.006 ± 0.003, p = 0.011; n = 209). In addition, there was an increase in the beta band SPL between FEF LFPs and IT spikes during the Pref, In compared to the NPref, In condition, indicative of the object selectivity of this SPL (ΔSPL = 0.007 ± 0.002, p = 0.002, n = 228). In contrast, neither the SPL between FEF spikes and IT LFPs, nor within-IT SPL, was predictive of performance. There was no difference in the beta band SPL between FEF spikes and IT LFPs for correct vs. wrong trials (ΔSPL = 0.002 ± 0.004, p = 0.693; n = 96; Fig. 3c). The beta band SPL between IT LFPs and IT spikes was not significantly different for correct vs. wrong trials (ΔSPL = 0.004 ± 0.004, p = 0.077, n = 140 neuron- LFP pairs; Fig. 3d). Thus, inter-areal, but not within area, beta coupling (PPL and SPL) predicted memory performance.

### Beta coupling between the FEF and IT as a predictor of IT’s object discriminability

Object-selective persistent activity in IT has often been suggested as a basis of WM maintenance^15^. Beta coupling can also reflect the identity of the object held in WM^9,16^. We sought to determine if there was a relationship between beta coupling and the object-selective persistent activity in IT. To test for this relationship, we quantified beta coupling on individual trials (see Methods). We sorted trials by beta band PPL value, and designated the third of trials with the highest PPL as “High” and the lowest third as “Low” trials (Methods). Using ROC analysis, we calculated the time-course of the object discriminability of IT spiking activity for High and Low PPL trials (Fig. 4a). For the In condition, High PPL trials had more object discriminability than Low PPL trials during the delay period (ΔAUC = 0.040 ± 0.016, p =0.021 n = 90; Fig. 4b). For the Out condition, there was no difference in object discriminability between High and Low PPL trials (ΔAUC = −0.025 ± 0.015, p = 0.209, n = 90, Fig. S2). Figure 4c shows the modulation index (MI) of object discriminability values (see Methods) for High vs. Low PPL trials for object-selective IT units, across time for the In (red) and Out (blue) conditions. There was an increase in the modulation of object discriminability by PPL during the delay period for the In condition compared to the Out condition (ΔMI = 0.064 ± 0.019, p = 0.003, n = 90; Fig. 4d). We next sought to test whether this modulation by beta coupling was dependent on the strength of object discriminability of the IT unit. Across the IT population, for the In condition, there was a correlation between IT units’ object discriminability and the discriminability difference for High vs. Low PPL trials (r = 0.174, p < 10^-6^, n = 146; Fig. 4e). No such correlation was observed for the Out condition (r = 0.056, p = 0.167, n = 146; Fig. 4f). These results support the idea that beta coupling between the FEF and IT predicts the maintenance of object identity within IT cortex.

**Figure 4.**
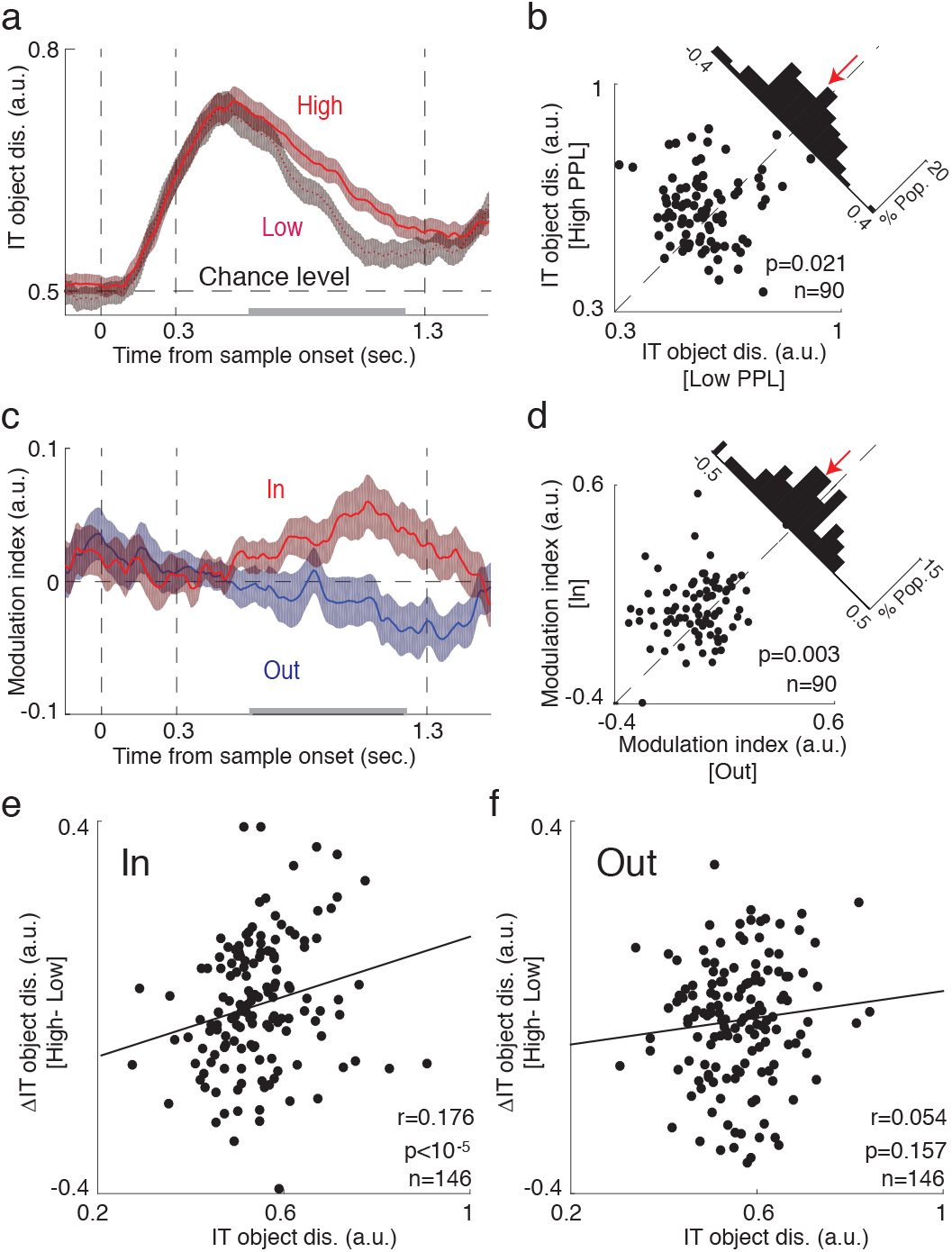
Enhancement of object coding by IT spiking activity during high beta PPL. **a,** Object discriminability was greater on trials with high beta PPL. Time-course of the object discriminability values for High (light red) vs. Low (dark red) PPL trials for object-selective IT units. Data were smoothed within a window of 1 ms and represented as mean ± SE. Gray bar indicates portion of delay period used for analysis in (b). **b,** Scatterplot illustrates object discriminability during the delay period for High vs. Low beta PPL trials, for the In condition. Histograms in the upper right show the difference in discriminability between High and Low PPL trials. **c,** Plot shows the time-course of the modulation index of object discriminability (High vs. Low PPL trials), for In (red) and Out (blue) conditions. Data were smoothed with a window of 1ms and represented as mean ± SE. **d,** Scatterplot illustrates the modulation index of object discriminability (High vs. Low PPL) during the delay period forIn vs. Out conditions. Histograms in the upper right show the difference between In and Out. **e,** Scatter plot shows the correlation between discriminability for each IT unit (x-axis) and the difference in discriminability for that unit between High and Low beta PPL trials (y-axis). **f,** No correlation between IT units’ discriminability and the discriminability difference for High vs. Low beta PPL trials for the Out condition. Same as (e) but for the Out condition.

## Discussion

Prefrontal areas are known to play an important role in WM. In order to understand how PFC interacts with sensory areas during WM, we simultaneously recorded from a prefrontal area (FEF) where neurons exhibit spatially-selective persistent activity and from IT, where neurons exhibit object-selective persistent activity, during an object WM task (Fig. 1). Whereas neither the average spiking activity nor the LFPs within either of these areas strongly predicted behavioral performance (Fig. 1c-d), the phase-phase locking between the two areas, specifically in the beta frequency range, was robustly correlated with the behavioral outcome (Fig. 2a-c). This coordination between areas was selective for both the spatial and object content of WM (Fig. 2d-e), and FEF oscillations led those in IT (Fig. 3a) and determined the timing of its spiking activity (Fig. 3b). This inter-areal oscillatory coherence not only predicted the behavioral outcome, it also predicted the strength of object discriminability within IT during memory maintenance (Fig. 4).

Our results support the idea that the interaction between prefrontal and sensory areas contributes to WM maintenance. Although we cannot rule out the possibility that one or more other areas drive WM selective signals in both the FEF and IT during WM, the findings that FEF leads IT in phase, and that inter-areal coherence predicts IT object selectivity, are consistent with a direct gating of object-selective persistent activity in IT by the FEF. This leading role of the FEF dovetails with previous results showing Granger-based causality of the FEF on IT during memory maintenance, but not the reverse^17^. The known influence of FEF neurons on the visual responses in visual cortex^18–20^, and the evidence that FEF neurons send memory-related spiking activity directly to visual areas^21^, further support the idea that FEF spiking activity has a direct impact on IT during WM. It is important to note that this is not the first time a spatial signal has been shown to enhance feature selectivity. In the case of covert spatial attention^22,23^, spatial cues have been shown to increase feature information in V4^24,25^ and IT^26^. A similar gating of feature information by a spatial signal is observed when attention is exerted overtly, when the representation of targets of upcoming eye movements are enhanced^27–30^. More specifically, FEF spiking activity, lacking robust feature selectivity, has been shown to contribute to feature selectivity in these areas^18–20,31,32^. A spatial tag has also been suggested as a basis for binding object features^33,34^, including during short term memory^35,36^; consistent with this hypothesis, retroactive spatial cues can improve object memory^37–41^. In our data, inter-areal coherence with the FEF could reflect the deployment of spatial attention for feature binding. However, in the absence of a competing stimulus, this FEF contribution might not be apparent^42^. These results suggest that perhaps, similar to the case of spatial attention, a non-selective signal highlighting relevant information is sufficient to boost object-selective spiking activity in IT.

The coordination between the FEF and IT occurs mostly within the beta band. In prefrontal and visual areas, increased beta power has been reported during the delay period of a variety of tasks^9,43–49^, leading to the suggestion that maintenance of the current state, whether motor or cognitive, is the unifying principle of beta oscillations^50^. Indeed, beta band coherence between IT sites is correlated with object WM performance in humans^48^. Such a content-agnostic role for beta oscillations in maintaining current states and information, whatever those might be, is consistent with the idea of a nonselective gain signal from the FEF gating object information within sensory areas. Note that despite the lack of feature selectivity in the FEF, we nevertheless observed spatial and feature selective *coordination* between the FEF and IT (Fig. 3d-e and Fig. S1). The finding that an area without feature selectivity can develop a content-specific coordination with a feature selective sensory area is further evidence that nonselective signals can nevertheless gate both selective sensory and memory spiking activity.

## Methods

### Animals and recordings

Two adult male rhesus monkeys (10 and 11 kg) were used in this study. M1’s experiments were performed in the School of Cognitive Science at IPM, and M2’s experiments were performed at Stanford University. All experimental procedures were in accordance with the National Institutes of Health Guide for the Care and Use of Laboratory Animals and the Society for Neuroscience Guidelines and Policies. The protocols for all experimental, surgical, and behavioral procedures were approved by the Institute Fundamental Science committee for monkey 1, and by the Stanford University Institutional Animal Care and Use Committee for monkey 2. Structural magnetic resonance imaging (M1, M2) and CT scan (M1) were performed to locate the arcuate sulcus and prelunate gyrus for the placement of a recording chamber in a subsequent surgery. All surgical procedures were carried out under Isoflurane anesthesia and strict aseptic conditions. Prior to undergoing behavioral training, each animal was implanted with a stainless-steel custom-made chamber, attached to the skull using orthopedic titanium screws and dental acrylic. For M1, within the chamber 30×70 mm craniotomies was performed (craniotomy was 5 mm to 30 mm A/P, and 0 mm to 23 mm M/L). For M2, within the chamber a 20mm diameter craniotomy was performed (chambers were started at 25 mm A/P, 23 mm M/L and ended 5 mm A/P, 23 mm M/L).

### Behavioral monitoring

Animals were seated in custom-made primate chairs, with their head restrained and a tube to deliver juice rewards placed in their mouth. For M1, eye position was monitored with an infrared optical eye tracking system (EyeLink 1000 Plus Eye Tracker, SR Research Ltd, Ottawa CA), with a resolution of < 0.01° RMS; eye position was monitored and stored at 2 KHz. The EyeLink PM-910 Illuminator Module and EyeLink 1000 Plus Camera (SR Research Ltd, Ottawa CA) were mounted in front of the monkey, and captured eye movements. Stimulus presentation and juice delivery were controlled using custom software, written in MATLAB using the MonkeyLogic toolbox (Asaad et al., 2013). Visual stimuli were presented on an LED-lit monitor (Asus VG248QE: 24in, resolution 1920×1080, 144 Hz refresh rate), positioned 65.5 cm in front of the animal’s eyes. A photodiode (OSRAM Opto Semiconductors, Sunnyvale CA) was used to record the actual time of stimulus appearance on the monitor. For M2, eye position was monitored with a scleral search coil and digitized at 500 Hz (CNC Engineering, Seattle, WA). The spatial resolution of eye position measurements was << 0.1 deg. Stimulus presentation, data acquisition and behavioral monitoring were controlled by CORTEX system. Visual stimuli were presented on a 29° x 39° colorimetrically calibrated CRT monitor (Mitsubishi Diamond Pro 2070SB-BK) with medium short persistence phosphors. A photodiode (OSRAM Opto Semiconductors, Sunnyvale CA) was used to record the actual time of stimulus appearance on the monitor, with a continuous signal sampled and stored at 32 KHz.

### Behavioral tasks

#### Eye calibration

The fixation point, a ~0.25 dva white circle, appeared either centrally or displaced by +- 10dva in the horizontal or vertical axis, and the monkey maintained fixation within a ± 1.5 dva window for 1.5 s to receive a reward.

#### FEF RF estimation

We identified FEF sites by eliciting short-latency, fixed-vector saccadic eye movements with trains (50-100 ms) of biphasic current pulses (#50 mA; 250 Hz; 0.25 ms duration). RF mapping was conducted by having the monkey fixate within a ± 1.5 dva window around the fixation point (central or displaced by +-10 dva), while stimulation was applied to evoked a saccade. The average endpoint of evoked saccades (landing points) was used to estimate the center of the FEF RF.

#### Finding preferred stimuli of IT units

Rapid serial visual presentation (RSVP) task was used to quickly screen 44 sample objects and identify at least one which evoked IT visual responses for subsequent use in the DMS task. Monkeys fixated within a ±1.5 dva window around the central fixation point, and 10 objects (from 44 possible) were pseudo-randomly presented for 150ms each with 100ms between stimuli. Responses from the recording site in IT cortex were monitored audibly and visually by the experimenter. Stimuli evoking the highest response (‘Pref’), an intermediate response (‘Inter), and little or no response (‘NPref’) were selected for use in subsequent behavioral task.

#### Object delayed-match-to-sample task

The DMS task is a widely used WM task; the version we use requires object WM. Monkeys fixated within a ± 1.5 dva window around the central fixation point. After 500 ms of fixation, a 3 dva sample object was presented either inside the FEF RF, or 180 degrees away in the opposite hemifield, and remained onscreen for 300 ms. The animal then remembered the sample identities while maintaining fixation for 1 s (memory period) before the central fixation point was removed and two targets appeared (the sample object and a distractor). Target positions were rotated 90 degrees relative to the sample location so no target appeared at the sample location, nor did the sample location predict the correct target location. The monkey had to shift its gaze to a ± 4 dva window around the sample object to receive a juice reward. If the monkey completes the trial but selects the distractor object, it was labeled a wrong trial. Note that in this version of the task, the position of the sample is irrelevant for task performance.

#### Neurophysiological recording

Two electrodes were mounted on the recording chamber and positioned within the craniotomy area using two Narishige two-axis platforms allowing continuous adjustment of the electrodes’ position. Two 28- gauge guide tubes were lowered to contact or just penetrate the dura, using a manual oil hydraulic micromanipulator (Narishige, Tokyo, Japan). For M1 and FEF of M2, varnish-coated tungsten microelectrodes (FHC, Bowdoinham, ME), shank diameter 200-250 μm, impedance 0.2–1 MΩ (measured at 1 kHz), were advanced into the brain for the extracellular recording of neuronal activity. For IT recordings in M2, a 32 gauge (235mm outer diameter) guide tube was advanced ~15mm through the brain (at a rate of ~.75mm/minute) by a custom-modified electronic motor-driven microdrive, stopping at or just above the upper bank of the Superior Temporal Sulcus. An electrode (75-100 *μ* m diameter) was then advanced another 5-12mm using a hydraulic microdrive (Narashige). Activity was recorded extracellularly with varnish-coated tungsten microelectrodes (FHC) of 0.2–1.0 M Ω impedance (measured at 1 kHz). For M1, single-electrode recordings used a RESANA pre-amplifier and RESANA amplifier, filtering from 300 Hz – 5 KHz for spikes and 0.1 Hz – 9 KHz for LFPs. Spike waveforms and continuous data were digitized and stored at 40 kHz for offline spike sorting and data analysis. For M2, extracellular waveforms were digitized and classified as single neurons using online template-matching and window-discrimination techniques (FHC Inc., Bowdoinham, ME). Spike waveforms were sorted manually by Plexon offline sorting. Area IT was identified based on stereotaxic location, position relative to nearby sulci, patterns of gray and white matter, and response properties of units encountered. Area FEF was identified based on stereotaxic location, position relative to acute sulcus, patterns of gray and white matter, and also microstimulation. By single-electrode exploration and electrical stimulation, areas posterior of the arcuate sulcus (evoking movements of hand and face) and the FEF (evoking saccadic eye movements) were identified within the recording chamber prior to beginning data collection.

Both single and multiunit data were included in all analysis. Data are reported from 58 IT sites (36 M1, 22 M2) and 86 FEF sites (51 M1, 35 M2). LFP data of 28 IT sites (14 M1, 14 M2) was discarded prior to any analysis due to noise or data acquisition issues. When comparing correct and wrong trials, certain sites and units were excluded due to low numbers of wrong trials (< 3 wrong trials); number of sites and units included in each analysis are included in the text with the statistics. For IT LFPs, object selectivity of the site was determined based on local multiunit responses (similar to ^51^); in figure 2d and figure S1c, LFPs from 7 IT sites were excluded from analysis because multiunit activity was not object selective.

## Data analysis

All analyses were carried out with custom code written in Matlab (MathWorks). The raw LFP data was low-pass filtered (1-300 Hz) and resampled at 1 kHz. The LFP for each site was normalized by subtracting the mean and dividing by the s.d. (across all timepoints and trials). Analysis focused on the 600 ms window in the late delay (300-900 ms after stimulus offset), in order to exclude both the stimulus evoked response and presaccadic activity (~100 ms before target onset); this is the time window used for analysis unless otherwise stated. In the trial-matching procedure, the number of correct and wrong trials was matched by subsampling the larger distribution 1001 times, and taking the mean of that distribution.

For firing rate calculations (Fig. 1c), individual spike trains were smoothed with a 10 ms gaussian kernel, and the mean response was calculated across trials for each condition. The normalized firing rate was calculated according to the formula NFR(t) =[Rate(t) – Baseline] / (MaxRate – Baseline), where Baseline was the average firing rate during the 500 ms of the fixation period before sample onset, and MaxRate was the maximum during the task.

To calculate LFP power and phase (Fig. 1d), we extracted the instantaneous amplitude and analytical phase as a function of time and frequency by convolving the raw real-valued time series x(t) with the complex Morlet wavelet w(t, f_0_) to obtain the complex output signal y(t, f_0_), also denoted as the analytic signal, where f_0_ denotes the desired center frequency of the wavelet function. We used a value of c = 7 wavelet oscillations. The center frequencies f_0_ were 1-130 Hz.

Time-frequency composition of the power spectrum was calculated by analytical amplitude of wavelet for each area. We averaged power values for each condition. In order to remove the 1/f effect we calculated Z-transformed values for each trial by subtracting the mean of the baseline and dividing by the standard deviation of the baseline across sites for each condition. The time frequency map was smoothed with a window of 2 Hz in frequency and 100 ms in time. Then, Z-transformed values were averaged across sites.

The power-power correlation was calculated over time and frequency. For each point (f.t) the Kendall correlation between FEF power and IT power across trials was calculated for each condition. In order to compare the conditions, we calculated Z-transformed correlation values by subtracting the value during the baseline and dividing by the standard deviation across pairs for each condition.

In order to quantify within area PLV (Table S1), we considered the phase value of each FEF and IT LFP using analytical phase calculated by wavelet. We calculated PLV for each time and frequency according to equation 1

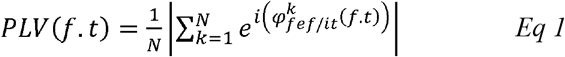

where 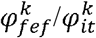 were the instantaneous phase values in trail k-th in frequency f and time bin t, and N was number of trials. The PLV was in [0 1], where 1 indicates high phase locking and 0 is no phase locking. In order to compare the conditions, we calculated Z-transformed PLV values by subtracting its value during baseline and dividing by the standard deviation across pairs for each condition.

In order to quantify PPL between areas (Fig. 2, Fig. S1), we considered the phase difference between the FEF and IT using analytical phase calculated by wavelet. We calculated PPL for each time and frequency according to equation 2

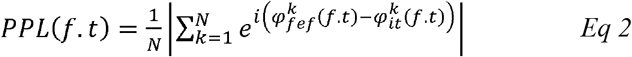

where 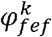 and 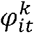 were the instantaneous phase values in trail k-th in frequency f and time bin t, and N was number of trials. The PPL was in [0 1], where 1 indicates high phase locking and 0 is no phase locking. Although within-area phase locking did not significantly differ between correct and wrong trials in the beta band (see Supplementary Information), we nevertheless used a trial shuffling procedure to remove any effect of within-area phase locking from the inter-area phase locking calculation. A shuffled distribution for each pair was calculated by randomly shuffling the pairing of trials for each site 1001 times. We then subtracted the mean of the shuffled distribution from the raw PPL value. In order to compare the conditions, we calculated Z-transformed PPL values by subtracting its value during baseline and dividing by the standard deviation across pairs for each condition. The time frequency map was smoothed with a window of 2 Hz in frequency and 100 ms in time. Then, Z-transformed values were averaged across sites. PPL values in figure 2c-e were averaged across the delay period for the beta band.

The phase difference distributions were calculated similar to PPL, equation 2, except instead of absolute value the angle was calculated. For each pair the circular average was calculated over beta band frequencies and the delay period analysis window. The circular histogram of correct and wrong trials for Pref, In condition was plotted in figure 3a. Von Mises circularity tests were used to check the concentration of the distributions of the phase difference values of pairs. Concentration parameter, κ values, between the correct and wrong trials were compared using a bootstrap test (n = 1001).

SPL measurements in figure 3 were performed on 209 spike-LFP pairs (IT units, FEF LFP), 96 spike-LFP pairs (FEF units, IT LFP), and 140 spike-LFP pairs (IT units, IT LFP) in both monkeys. The LFP for each trial was normalized by subtracting its mean and dividing by its standard deviation. We used a wavelet transform to calculate the analytic signal of all components. Instantaneous phases of each component were quantified by calculating the angles corresponding to the analytic signal. For each spike, we made a vector with the instantaneous phase at the spike time and an amplitude of one. The spikes from all trials were pooled together, and then we used the vector averaging method to calculate the SPL magnitude. In order to control for any effects of different numbers of spikes and trials between the two conditions, we considered a fixed window of 15 spikes and measured the SPL magnitude for the spikes of each window, and then took the mean across these windows^46^.

In figure 4, PPL was calculated for single trials to split data into High and Low PPL trials. Single-trial PPL was calculated based on equation 2 across the delay window and baseline in one trial^52^. N was the number of time points in the window. We subtracted the PPL value of the baseline from the delay period PPL to remove the 1/f effect. Trials were ranked based on single trial PPL values within each condition. We selected trials with PPL< 33th percentile as Low and trials with PPL> 66th percentile as High trials. Object discriminability was quantified using the receiver operating characteristic (ROC) analysis^53^. ROC analysis was performed on the distributions of neuronal firing rate. The areas under ROC curves (AUC) when comparing responses to different sample objects were used as an index of object discriminability, and were calculated as in previous studies^31^. For the timecourse of object discriminability of IT units, ROC was calculated for a jumping 300 ms window (10 ms steps). ROC values were calculated with a minimum of 3 trials per condition. For figure 4a and figure S2a, we selected IT units with a discriminability of AUC >=0.51. For figure 4c-d, we calculated the object AUC during the delay period for all units for High and Low PPL trials, then defined the modulation index as ((AUC_High_ – AUC_Low_)/ (AUC_High_ + AUC_Low_). For the scatter plots in 4b and 4d, we averaged the values over two adjacent 300ms windows within the delay period. In figure 4e-f, we calculated the Kendal correlation between the PPL difference (High – Low trials) and overall object discriminability, across sites. P-value was calculated by permutation test (n = 1001).

## Statistical analysis

Statistical analyses were performed using the Wilcoxon’s signed-rank test (for paired comparisons) or rank sum (for unpaired comparisons), unless otherwise specified. All p-values and effect sizes are reported up to 3 digits and values below 0.001 are reported as P < 10^-n^.

## References

1. Miller, B. T. & D’Esposito, M. Searching for “the top” in top-down control. Neuron 48, 535–538 (2005).

2. Fuster, J. M. & Alexander, G. E. Neuron activity related to short-term memory. Science (80-.). 173, 652–654 (1971).

3. Lara, A. H. & Wallis, J. D. The role of prefrontal cortex in working memory: a mini review. Front. Syst. Neurosci. 9, 173 (2015).

4. Harrison, S. A. & Tong, F. Decoding reveals the contents of visual working memory in early visual areas. Nature 458, 632–635 (2009).

5. O Scalaidhe, S. P., Wilson, F. A. & Goldman-Rakic, P. S. Areal segregation of face-processing neurons in prefrontal cortex. Science (80-.). 278, 1135–8 (1997).

6. Goldman-Rakic, P. S. The prefrontal landscape: implications of functional architecture for understanding human mentation and the central executive. Philos. Trans. R. Soc. London. Ser. B Biol. Sci. 351, 1445–1453 (1996).

7. Christophel, T. B., Klink, P. C., Spitzer, B., Roelfsema, P. R. & Haynes, J.-D. The distributed nature of working memory. Trends Cogn. Sci. 21, 111–124 (2017).

8. Serences, J. T. Neural mechanisms of information storage in visual short-term memory. Vision Res. 128, 53–67 (2016).

9. Salazar, R. F., Dotson, N. M., Bressler, S. L. & Gray, C. M. Content-specific fronto-parietal synchronization during visual working memory. Science (80-.). 437, 1097–1101 (2012).

10. Mendoza-Halliday, D., Torres, S. & Martinez-Trujillo, J. C. Sharp emergence of feature-selective sustained activity along the dorsal visual pathway. Nat. Neurosci. 17, 1255–1262 (2014).

11. Supèr, H., Spekreijse, H. & Lamme, V. A. F. A neural correlate of working memory in the monkey primary visual cortex. Science (80-.). 293, 120–124 (2001).

12. Shahidi, N., Andrei, A. R., Hu, M. & Dragoi, V. High-order coordination of cortical spiking activity modulates perceptual accuracy. Nat. Neurosci. 22, 1148 (2019).

13. Zaksas, D. & Pasternak, T. Directional signals in the prefrontal cortex and in area MT during a working memory for visual motion task. J. Neurosci. 26, 11726–11742 (2006).

14. Clark, K. L., Noudoost, B. & Moore, T. Persistent spatial information in the frontal eye field during object-based short-term memory. J. Neurosci. 32, 10907–10914 (2012).

15. Fuster, J. M. & Jervey, J. P. Inferotemporal neurons distinguish and retain behaviorally relevant features of visual stimuli. Science (80-.). 212, 952–955 (1981).

16. Lee, H., Simpson, G. V, Logothetis, N. K. & Rainer, G. Phase locking of single neuron activity to theta oscillations during working memory in monkey extrastriate visual cortex. Neuron 45, 147–56 (2005).

17. Hu, M. et al. Copula regression analysis of simultaneously recorded frontal eye field and inferotemporal spiking activity during object-based working memory. J. Neurosci. 35, 8745–57 (2015).

18. Noudoost, B. & Moore, T. Control of visual cortical signals by prefrontal dopamine. Nature 474, 372–375 (2011).

19. Monosov, I. E. & Thompson, K. G. Frontal eye field activity enhances object identification during covert visual search. J. Neurophysiol. 102, 3656–3672 (2009).

20. Moore, T. & Armstrong, K. M. Selective gating of visual signals by microstimulation of frontal cortex. Nature 421, 370–373 (2003).

21. Merrikhi, Y. et al. Spatial working memory alters the efficacy of input to visual cortex. Nat. Commun. 8, 15041 (2017).

22. Noudoost, B., Chang, M. H., Steinmetz, N. A. & Moore, T. Top-down control of visual attention. Curr. Opin. Neurobiol. 20, 183–90 (2010).

23. Reynolds, J. H. & Chelazzi, L. Attentional modulation of visual processing. Annu. Rev. Neurosci. 27, 611–47 (2004).

24. McAdams, C. J. & Maunsell, J. H. R. Effects of attention on orientation-tuning functions of single neurons in macaque cortical area V4. J. Neurosci. 19, 431–441 (1999).

25. Reynolds, J. H., Pasternak, T. & Desimone, R. Attention increases sensitivity of V4 neurons. Neuron 26, 703–714 (2000).

26. Zhang, Y. et al. Object decoding with attention in inferior temporal cortex. Proc. Natl. Acad. Sci. 108, 8850–5 (2011).

27. Moore, T. & Chang, M. H. Presaccadic discrimination of receptive field stimuli by area V4 neurons. Vision Res. 49, 1227–1232 (2009).

28. Moore, T. Shape representations and visual guidance of saccadic eye movements. Science (80-.). 285, 1914–7 (1999).

29. Han, X., Xian, S. X. & Moore, T. Dynamic sensitivity of area V4 neurons during saccade preparation. Proc. Natl. Acad. Sci. 106, 13046–13051 (2009).

30. Moore, T., Tolias, A. S. & Schiller, P. H. Visual representations during saccadic eye movements. Proc. Natl. Acad. Sci. 95, 8981–8984 (1998).

31. Armstrong, K. M. & Moore, T. Rapid enhancement of visual cortical response discriminability by microstimulation of the frontal eye field. Proc. Natl. Acad. Sci. 104, 9499–9504 (2007).

32. Noudoost, B., Clark, K. L. & Moore, T. A distinct contribution of the frontal eye field to the visual representation of saccadic targets. J. Neurosci. 34, 3687–3698 (2014).

33. Treisman, A. & Schmidt, H. Illusory conjunctions in the perception of objects. Cogn. Psychol. 14, 107–141 (1982).

34. Treisman, A. M. & Gelade, G. A feature-integration theory of attention. Cogn. Psychol. 12, 97–136 (1980).

35. Treisman, A. & Zhang, W. Location and binding in visual working memory. Mem. Cognit. 34, 1704–1719 (2006).

36. Wheeler, M. E. & Treisman, A. M. Binding in short-term visual memory. J. Exp. Psychol. Gen. 131, 48–64 (2002).

37. Nobre, A. C., Griffin, I. C. & Rao, A. Spatial attention can bias search in visual short-term memory. Front. Hum. Neurosci. 1, 4 (2007).

38. Lepsien, J., Griffin, I. C., Devlin, J. T. & Nobre, A. C. Directing spatial attention in mental representations: Interactions between attentional orienting and working-memory load. Neuroimage 26, 733–43 (2005).

39. Griffin, I. C. & Nobre, A. C. Orienting attention to locations in internal representations. J. Cogn. Neurosci. 15, 1176–1194 (2003).

40. Matsukura, M., Luck, S. J. & Vecera, S. P. Attention effects during visual short-term memory maintenance: Protection or prioritization? Percept. Psychophys. 69, 1422–1434 (2007).

41. Makovski, T., Sussman, R. & Jiang, Y. V. Orienting attention in visual working memory reduces interference from memory probes. J. Exp. Psychol. Learn. Mem. Cogn. 34, 369–80 (2008).

42. Clark, K. L., Noudoost, B. & Moore, T. Persistent spatial information in the FEF during object-based short-term memory does not contribute to task performance. J. Cogn. Neurosci. 26, 1292–1299 (2014).

43. Lundqvist, M. et al. Gamma and Beta Bursts Underlie Working Memory. Neuron 90, 152–164 (2016).

44. Tallon-Baudry, C., Bertrand, O. & Fischer, C. Oscillatory synchrony between human extrastriate areas during visual short-term memory maintenance. J. Neurosci. 21, 1–5 (2001).

45. Siegel, M., Warden, M. R. & Miller, E. K. Phase-dependent neuronal coding of objects in shortterm memory. Proc. Natl. Acad. Sci. 106, 21341–21346 (2009).

46. Bahmani, Z., Daliri, M. R., Merrikhi, Y., Clark, K. & Noudoost, B. Working Memory Enhances Cortical Representations via Spatially Specific Coordination of Spike Times. Neuron 97, 967–979.e6 (2018).

47. Lundqvist, M., Herman, P., Warden, M. R., Brincat, S. L. & Miller, E. K. Gamma and beta bursts during working memory readout suggest roles in its volitional control. Nat. Commun. 9, 1–12 (2018).

48. Tallon-Baudry, C., Mandon, S., Freiwald, W. A. & Kreiter, A. K. Oscillatory synchrony in the monkey temporal lobe correlates with performance in a visual short-term memory task. Cereb. Cortex 14, 713–720 (2004).

49. Deiber, M. P. et al. Distinction between perceptual and attentional processing in working memory tasks: A study of phase-locked and induced oscillatory brain dynamics. J. Cogn. Neurosci. 19, 158–172 (2007).

50. Engel, A. K. & Fries, P. Beta-band oscillations-signalling the status quo? Curr. Opin. Neurobiol. 20, 156–165 (2010).

## Method References

51. Hung, C. P., Kreiman, G., Poggio, T. & DiCarlo, J. J. Fast readout of object identity from macaque inferior temporal cortex. Science (80-.). 310, 863–866 (2005).

52. Lachaux, J.-P. et al. Studying single-trials of phase synchronous activity in the brain. Int. J. Bifurc. Chaos 10, 2429–2439 (2000).

53. Green, D. M., Swets, J. A. & others. Signal detection theory and psychophysics. 1, (Wiley New York, 1966).

