## Supplementary Information for "Frontotemporal Coordination Predicts Working Memory Performance and its Local Neural Signatures"

### Supplementary Results

#### Spatial selectivity of neural activity in the FEF:

During the delay period, there was an increase in FEF gamma band power ($\Delta$power = 0.256 ± 0.062, $p < 10^{-5}$, n = 86 sites) and a decrease in FEF beta band power compared to baseline ($\Delta$power = -0.488 ± 0.044, $p < 10^{-14}$, n = 86 sites; Fig. 1d top). Power during the delay period also differed based on sample location for both the beta band ($\Delta$power = 0.106 ± 0.032, $p < 10^{-4}$, n= 86 sites) and the gamma band ($\Delta$power = 0.253 ± 0.062, $p < 10^{-7}$, n = 86 sites).

#### Object selectivity of neural activity in IT:

The IT population did not exhibit a significant increase in spiking activity during the delay period compared to baseline for the preferred object ($\Delta$NFR = -0.004 ± 0.006, p = 0.930, n = 228), but did show a significant decrease in spiking activity for the non-preferred object ($\Delta$NFR = -0.049 ± 0.005, $p < 10^{-15}$, n = 228).

In IT, there was an increase in delay period alpha band power compared to baseline ($\Delta$power = 0.056 ± 0.067, p = 0.031, n = 58 sites). Alpha-band IT power during the delay period differed for the preferred vs. non-preferred stimulus ($\Delta$power = 0.080 ± 0.029, p = 0.002, n = 58 sites).

#### Phase locking to stimulus onset:

LFP phase has been shown to lock to stimulus onset[14], and so we measured the phase-locking value (PLV) of LFP oscillations to the sample onset across trials, within FEF and IT. Within FEF, only the theta band ($\theta$, 4-8 Hz) PLV differed between the In and Out conditions ($\Delta PLV_{\theta}$ = -0.846 ± 0.077, $p < 10^{-12}$, n = 86) and there was a significant difference between theta PLV during correct vs. wrong trials ($\Delta PLV_{\theta}$ = -0.361 ± 0.059, $p < 10^{-7}$, n = 85 sites). Within IT, PLV was not object selective in any band, but $\theta$ and $\alpha$ band PLV were lower on correct trials compared to wrong trials ($\Delta PLV_{\theta}$ = -0.251 ± 0.073, p = 0.001, $\Delta PLV_{\alpha}$ = -0.211 ± 0.100, p = 0.011, n = 54 sites).

#### PPL for trial matching and without shuffle correction:

In order to control for differences in the number of trials, we repeated the PPL calculation using a trial matching procedure (Methods, Fig. S1a left), and found that in the beta band PPL was still greater for correct trials ($\Delta PPL_{Pref,In}$ = 0.393 ± 0.126, p = 0.002, n = 50). PPL statistics and data presented in figure 2 and the main text used a shuffling procedure to remove any effect of within-area phase locking (see Methods). Without this shuffling, there was still significantly higher beta band PPL on correct vs. wrong trials (Fig. S1a right; $\Delta PPL_{Pref,In}$ = 0.318 ± 0.097, p = 0.003, n = 50).

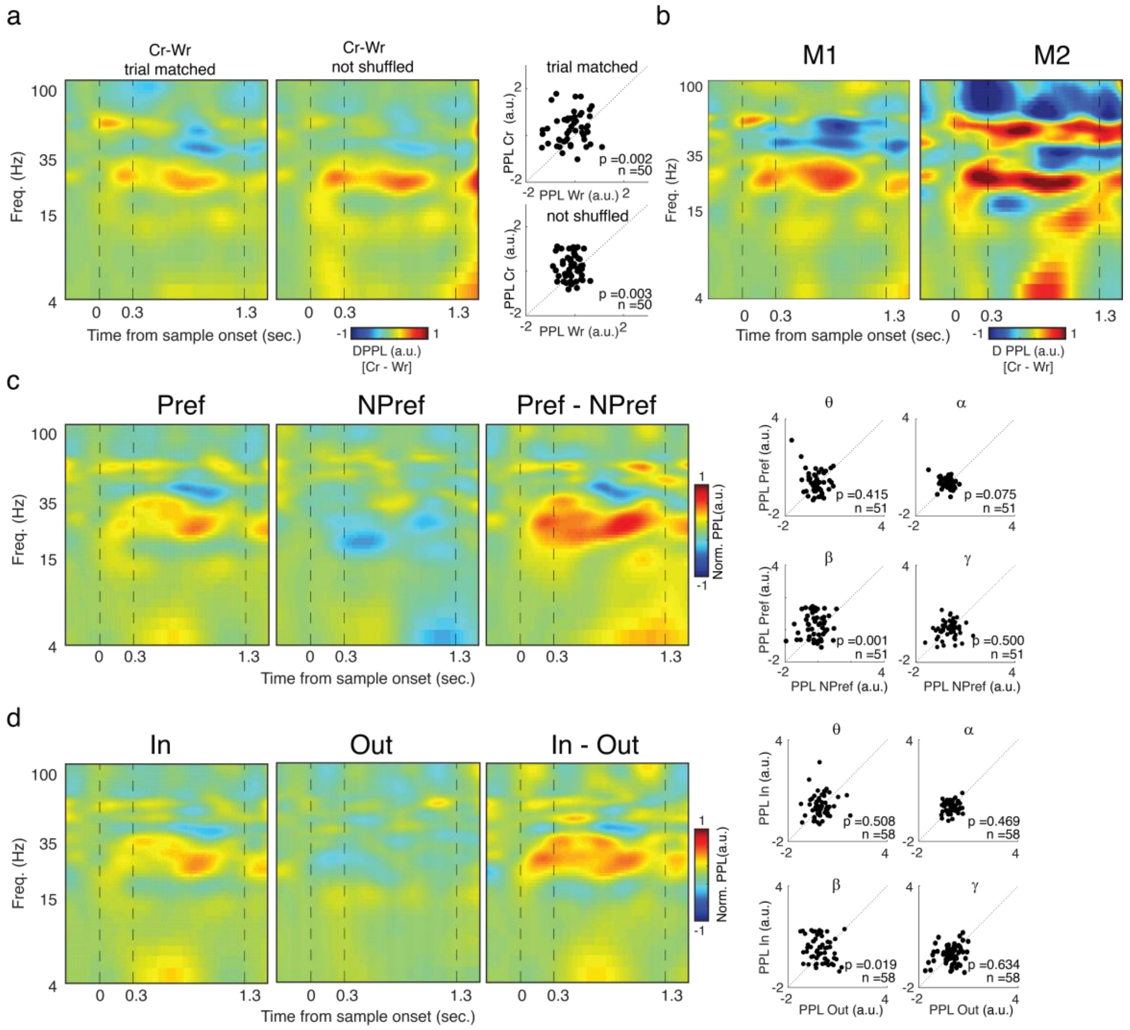


**Figure S1. PPL trial matching, M1 vs. M2, object and location coding.**

A, Beta band PPL was greater for correct trials when trial numbers were matched, and without the shuffle correction. Heatmaps show the time-frequency map of PPL, normalized to baseline across the course of the DMS task (n = 50 LFP recordings), for the IN condition, when number of correct and wrong trials were matched (left) and without shuffle correction (right). Scatterplots illustrate the average beta PPL for the correct vs. wrong trials with trial matching (top) and without shuffle correction (bottom). B, The trend toward higher beta band PPL on correct trials was present in both M1 and M2. Heatmaps show the time-frequency map of PPL difference between correct and wrong trials, across the course of the DMS task for the IN condition in M1 (left; n=35) and M2 (right; n=15). C, Inter-areal beta PPL encoded object identity during the delay period. Heatmap shows the time-frequency map of PPL, normalized to baseline across the course of the DMS task (n = 51 LFP pairs; correct trials, IN condition) for Pref (left) and NPref (middle) trials, and the difference between them (right). Scatter plots on the right show the mean delay-period PPL in different frequency bands for Npref (x-axis) vs. Pref (y-axis). D, Inter-areal beta PPL encoded the location of the sample during the delay period. Heatmap shows the time-frequency map of PPL, normalized to baseline across the course of the DMS task (n = 58 LFP pairs; correct trials, Pref condition) for In (left), Out (middle), and the difference between them (right). Scatter plots on the right show the mean delay-period PPL in different frequency bands for Out (x-axis) vs. In (y-axis).

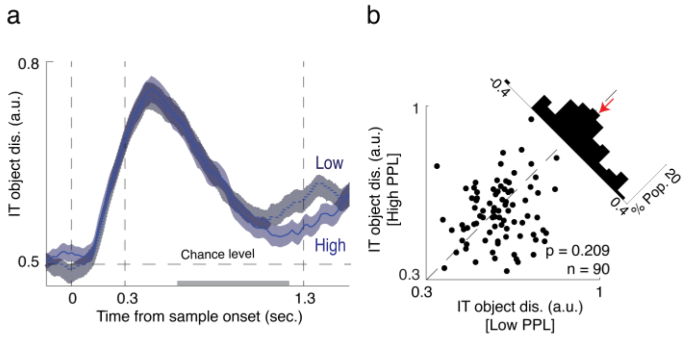


**Fig S2. Object coding in IT firing rate was enhanced for high beta PPL trials.**

A, Timecourse of the object discriminability values for High (green) vs. Low (blue) PPL trials for object-selective IT units, for the Out condition. Data were smoothed within a window of 1ms and represented as mean ± SE. Black bar indicates portion of delay period used for analysis in (B). B, Scatterplots illustrate object discriminability during the delay period for High vs. Low beta PPL trials, for the Out condition. Histogram in the upper right shows the difference between High and Low.

**Table S1, Statistics for relationship between within and inter-area measures and object, location, and performance.**

Neural measure is indicated by the label on the left (firing rate, LFP power or phase in a specific frequency band). Area (FEF, IT, or inter-area) is listed at the top. Selectivity measure (performance, Cr vs. Wr; location, In vs. Out; object, Pref vs. NPref) is indicated in the second row above the relevant column. Performance measures were calculated based on the Pref condition for IT, on the In condition for FEF and on the Pref, In condition for the interactions. Each box indicates the magnitude of the difference between conditions (Δ, mean ± SE), significance (p), and sample size (n) for the corresponding neural and selectivity measure. Color indicates significant increases (green) and decreases (gray) for the Cr, In, or Pref condition.

| | | *IT* | | *FEF* | | *FEF- IT Interaction* | | |
|---|---|---|---|---|---|---|---|---|
| | | Performance | Object | Performance | Location | Performance | Object | Location |
| *Firing rate* | | p = 0.282<br>Δ = 0.01± 0.00<br>n = 223 | $p < 10^{-15}$<br>Δ =0.05± 0.01<br>n = 228 | p =0.041<br>Δ = 0.01±0.00<br>n = 159 | $p < 10^{-7}$<br>Δ =0.06±0.01<br>n = 161 | | | |
| *LFP power* | θ | p = 0.997<br>Δ = -0.01± 0.06<br>n = 54 | p = 0.488<br>Δ = 0.08±0.06<br>n = 58 | p = 0.198<br>Δ =-0.04±0.04<br>n = 85 | $p < 10^{-7}$<br>Δ =-0.23±0.03<br>n = 86 | p = 0.540<br>Δ =-0.06±0.09<br>n = 50 | p = 0.757<br>Δ = 0.01±0.12<br>n = 51 | p = 0.257<br>Δ = 0.17±0.12<br>n = 58 |
| | *α* | p = 0.990<br>Δ = 0.01± 0.04<br>n = 54 | p = 0.002<br>Δ = 0.08±0.03<br>n = 58 | p = 0.373<br>Δ = 0.01±0.02<br>n = 85 | $p < 10^{-4}$<br>Δ =-0.02±0.03<br>n = 86 | p = 0.836<br>Δ = 0.05±0.10<br>n = 50 | p = 0.388<br>Δ =-0.05±0.09<br>n = 51 | p = 0.795<br>Δ = 0.04±0.07<br>n = 58 |
| | *β* | p = 0.265<br>Δ =-0.15±0.08<br>n = 54 | p = 0.679<br>Δ = 0.07±0.05<br>n = 58 | p = 0.078<br>Δ =-0.03±0.03<br>n = 85 | $p < 10^{-4}$<br>Δ =0.11± 0.03<br>n = 86 | p = 0.828<br>Δ = 0.06±0.14<br>n = 50 | p = 0.888<br>Δ =-0.02±0.15<br>n = 51 | p = 0.634<br>Δ = 0.02±0.12<br>n = 58 |
| | *γ* | p = 0.614<br>Δ =-0.03±0.02<br>n = 54 | p = 0.414<br>Δ = 0.00±0.02<br>n = 58 | p = 0.416<br>Δ =-0.01±0.03<br>n = 85 | $p < 10^{-7}$<br>Δ = 0.25±0.06<br>n = 86 | p = 0.229<br>Δ =-0.16±0.15<br>n = 50 | p = 0.339<br>Δ =-0.10±0.12<br>n = 51 | p = 0.217<br>Δ =-0.14±0.12<br>n = 58 |
| *LFP phase* | *θ* | p = 0.001<br>Δ =-0.25±0.07<br>n = 54 | p = 0.055<br>Δ =-0.10±0.07<br>n = 58 | $p < 10^{-7}$<br>Δ =-0.36±0.06<br>n = 85 | p < 0.001<br>Δ =-0.85±0.08<br>n = 86 | p = 0.896<br>Δ =0.07± 0.12<br>n = 50 | p = 0.414<br>Δ =0.18± 0.13<br>n = 51 | p = 0.508<br>Δ =0.10± 0.11<br>n = 58 |
| | *α* | p = 0.011<br>Δ =-0.21±0.10<br>n = 54 | p = 0.488<br>Δ =-0.04±0.09<br>n = 58 | $p < 10^{-6}$<br>Δ =-0.32±0.06<br>n = 85 | p = 0.309<br>Δ =-0.10±0.07<br>n = 86 | p = 0.117<br>Δ =0.12± 0.07<br>n = 50 | p = 0.075<br>Δ =0.13± 0.07<br>n = 51 | p = 0.469<br>Δ = 0.04±0.06<br>n = 58 |
| | *β* | p = 0.288<br>Δ =-0.17±0.14<br>n = 54 | p = 0.935<br>Δ =-0.06±0.12<br>n = 58 | P = 0.038<br>Δ =-0.16±0.08<br>n = 85 | p = 0.631<br>Δ =0.08± 0.12<br>n = 86 | p = 0.002<br>Δ = 0.44±0.13<br>n = 50 | p = 0.001<br>Δ = 0.52±0.14<br>n = 51 | p = 0.019<br>Δ = 0.36±0.15<br>n = 58 |
| | *γ* | p = 0.702<br>Δ = 0.01± 0.07<br>n = 54 | p = 0.427<br>Δ =-0.07±0.09<br>n = 58 | p = 0.010<br>Δ =-0.16±0.06<br>n = 85 | p = 0.798<br>Δ =-0.05±0.07<br>n = 86 | p = 0.121<br>Δ =-0.19±0.11<br>n = 50 | p = 0.500<br>Δ =0.03± 0.10<br>n = 51 | p = 0.634<br>Δ = 0.03±0.09<br>n = 58 |
